# Using predictive validity to compare associations between brain damage and behavior

**DOI:** 10.1101/2022.02.24.481802

**Authors:** John F. Magnotti, Jaclyn S. Patterson, Tatiana T. Schnur

## Abstract

Lesion-behavior mapping (LBM) provides a statistical map of the association between voxel-wise brain damage and individual differences in behavior. To understand whether two behaviors are mediated by damage to distinct regions, researchers often compare LBM weight outputs by either the Overlap method or the Correlation method. However, these methods lack statistical criteria to determine whether two LBM are distinct *vs*. the same and are disconnected from a major goal of LBMs: predicting behavior from brain damage. Without such criteria, researchers may draw conclusions from numeric differences between LBMs that are irrelevant to predicting behavior. We developed and validated a Predictive Validity Comparison method (PVC) that establishes a statistical criterion for comparing two LBMs using predictive accuracy: two LBMs are distinct if and only if they provide unique predictive power for the behaviors being assessed. We applied PVC to two lesion-behavior stroke data sets, demonstrating its utility for determining when behaviors arise from the same *vs*. different lesion patterns. Using region-of-interest based simulations derived from proportion damage from a large data set (*n* = 131), PVC accurately detected when behaviors were mediated by different regions (high sensitivity) vs. the same region (high specificity). Both the Overlap method and Correlation method performed poorly on the simulated data. By objectively determining whether two behavioral deficits can be explained by a single *vs*. distinct patterns of brain damage, PVC provides a critical advance in establishing the brain bases of behavior. We have developed and released a GUI-driven web app to encourage widespread adoption.

## Introduction

Lesion-behavior mapping is used to identify causal relationships between damage to brain regions and behavior (Bates et al., 2003; Rorden et al., 2007). A common application is to determine whether deficits in two behaviors are mediated by damage to the same or different brain areas. Although this type of comparison has been performed extensively in the literature (Alyahya et al., 2021; Arbula et al., 2020; Baldo et al., 2012; Barbey, Colom, Paul, et al., 2014; Barbey et al., 2013; Biesbroek et al., 2016; Binder et al., 2016; Chechlacz et al., 2013; Gläscher et al., 2009, 2010; Harvey & Schnur, 2015; Ivanova, Herron, Dronkers, & Baldo, 2021; Magnusdottir et al., 2013; Meyer et al., 2016; Mirman, Zhang, et al., 2015; Pini et al., 2021; Piras & Marangolo, 2009; Pirondini et al., 2022; Pustina et al., 2018; Rogalsky et al., 2015; Schwartz et al., 2009; Snider et al., 2020; Thye et al., 2021; Zhang et al., 2014) the field currently lacks a clear criterion for determining whether two behaviors require distinct lesion-behavior maps (LBMs) for maximum predictive accuracy from associated patterns of brain damage.

Identifying the brain regions necessary for human behavior is a well-established scientific endeavor beginning with single-case studies of stroke (e.g., Paul Broca’s report concerning patient Tan in 1861), subsequently extending to single case series (Dronkers, 1996; Schwartz & Dell, 2010), and finally gaining increased neural and statistical granularity with the advent of large sample, voxel-based lesion behavior mapping approaches (Bates et al., 2003; Rorden & Karnath, 2004). Typically, lesions are manually delineated on structural neuroimaging sequences slice-by-slice to create three-dimensional binary representations of the extent and location of brain damage after stroke at the level of the voxel (1-2mm cubed). These lesion masks are normalized to a common brain space to facilitate comparisons between individuals. Next, each voxel is assessed as to whether its damage is predictive of behavior, considered either independently in univariate approaches (Bates et al., 2003; Rorden et al., 2007) or jointly with other voxels as in multivariate approaches (Pustina et al., 2018; Zhang et al., 2014; cf. Ivanova et al., 2021).

The increasing availability of multimodal data has led recent research to focus more on the question of which brain map *types* (*e.g.,* lesion maps, functional connectivity, structural connectivity) best predict a single behavioral score (e.g., motor performance, language skills; Salvalaggio et al., 2020; Siddiqi et al., 2021; Siegel et al., 2016). Because these methods are largely concerned with determining the best-fitting brain map type (or combination of types) for a *single* behavior, researchers may only use these approaches to identify the brain regions necessary for a single behavior. However, if we wish to understand whether two behaviors depend on distinct brain regions, there are currently no statistically validated methods for making this determination.

When researchers compare LBMs, the most common approach is the Overlap method (Alyahya et al., 2021; Barbey, Colom, & Grafman, 2014; Biesbroek et al., 2016; Chechlacz et al., 2013; Ding et al., 2020; Gläscher et al., 2019; Harvey & Schnur, 2015; Magnusdottir et al., 2013; Meyer et al., 2016; Mirman, Chen, et al., 2015; Piras & Marangolo, 2007, 2009; Pustina et al., 2018; Snider et al., 2020; Thye et al., 2021; Thye & Mirman, 2018). In this approach, researchers compare LBMs depicting each behaviors’ necessary brain regions (commonly using t-values, beta weights, or voxel cluster size) by creating intersection and subtraction maps. Brain regions represented in one LBM but not the other are interpreted as distinctly involved in that particular behavior, whereas regional intersections are interpreted as regions critical to both behaviors. This technique, however, does not account for differences in LBMs caused by nuisance variance (e.g., sampling error, noise in the behavioral measure, etc.). Further, visual presentations of differences (or intersections) are rarely normalized or compared with a baseline, providing no information as to the scale of the differences, leading to the interpretation that any two LBMs that are not wholly identical are meaningfully different. These misuses and misinterpretations demonstrate that the Overlap method’s primary utility thus far has been as a qualitative visualization of the identical (the intersection map) and the non-identical (the subtraction map) areas of two LBMs.

In contrast, statistical measures are also used to compare LBMs. Some examples include determining the percent of overlapping voxels for each LBM (Binder et al., 2016; Gläscher et al., 2009), comparing the number of statistically significant voxels for each region implicated, and correlating LBM values (Ivanova et al., 2021; Pirondini et al., 2022; Schwartz et al., 2009; Thothathiri et al., 2012; Pini et al., 2022; Piras & Marangolo, 2009; Zhang et al., 2014). Although these methods involve statistical tests (Are the number of voxels in each map different? Is the correlation between LBM weights significant?), they fail to provide a usable threshold for determining LBM distinctness. For instance, finding that two LBMs are significantly correlated (*i.e., p* < 0.05) does not demonstrate that the LBMs are identical. Although the output of each method (percent overlap, LBM size, voxel-wise correlation) provides a measure of the degree of similarity between two LBMs, there is no clear way to convert these outputs to tests of the uniqueness of the underlying brain-behavior associations.

Here, we propose Predictive Validity Comparison (PVC) for testing the uniqueness of two LBMs fit to two behaviors. LBMs fitted to separate behaviors are declared “distinct” if and only if they provide unique predictive power for the behaviors being assessed. The critical step in PVC constructs two sets of predictions for individuals’ behaviors. The first set is generated under the null hypothesis that individual differences across the two behaviors are the result of a *single* lesion pattern. The second set is generated under the alternative hypothesis that individual differences across the two behaviors are the result of *distinct* lesion patterns. Only if the quality of the predictions under the alternative hypothesis is higher than under the null hypothesis do we conclude the LBMs are distinct. If the quality of the predictions under the null hypothesis are no worse than those from the alternative hypothesis, then we cannot conclude the LBMs are distinct. Instead, we conclude that either the behaviors are mediated by the same underlying lesion pattern, or more data are needed to distinguish their individual patterns.

We assessed the new method in three ways. First, we applied the PVC method to real data sets of different behaviors in stroke populations (Ding et al., 2020; Pustina et al., 2018) and compared the results with the Overlap and Correlation methods. Second, we assessed how the three methods were affected by changing parameters of the LBM fitting procedure. Finally, we assessed the ability of each method to correctly identify brain-behavior relationships in simulations for which the ground truth was known: the two behaviors resulted from damage to different brain regions versus the same brain region. To encourage adoption and extension of this method, we have released an open-source implementation of PVC complete with a user-friendly web-based interface.

## Method

The goal of the Predictive Validity Comparison (PVC) method is to determine whether individual differences across two behaviors are the result of a single pattern of lesion damage *vs*. distinct patterns of lesion damage. More concretely, can we accurately predict individuals’ behavioral scores using a single lesion-behavior map (LBM) or do we need a distinct LBM for each behavior? We first describe the two datasets used to assess the PVC method. Next, we provide the technical details of PVC. Then, we use the two datasets to compare the PVC method with two common LBM comparison methods: the Overlap method and the Correlation method. We compare the performance of each method across a range of hyperparameters used in fitting multivariate LBMs. Finally, we provide the results of a simulation analysis that assessed the sensitivity (accurately detecting when two behaviors have distinct neural bases) and specificity (accurately detecting when two behaviors have a shared neural basis) of all three methods when varying how strongly proportion damage in one region correlated with proportion damage in another region (the between-region lesion load correlation).

### Input data

#### The Moss Rehabilitation Research Institute (MRRI) Dataset

Lesion volumes and behavioral data for 131 patients (131 total lesion masks, but complete behavior was only available for 130 patients) were generously provided by the MRRI Neuro-Cognitive Rehabilitation Research Patient Registry (reported in previous publications, cf. Mirman et al., 2015; Pustina et al., 2016, 2018; Schwartz et al., 2009, 2011, 2012; Zhang et al., 2014). Patients had chronic left hemisphere stroke and clinical aphasia diagnoses (months post-onset: mean 44, range 2-387 months). All strokes were restricted to cortical infarcts in the left-hemisphere middle cerebral artery territory. Lesion maps were normalized to MNI space, pre-processed as described in Pustina et al. (2018), and available through the LESYMAP R package and ANTs registration (http://stnava.github.io/ANTs/; Avants et al., 2008; Avants, 2015). Behavioral scores included the Western Aphasia Battery Aphasia Quotient (WAB-AQ; Kertesz, 1982) and accuracy on the Philadelphia Naming Test (PNT; Roach et al., 1996) both reported in Pustina et al. (2018).

#### The Schnur Laboratory Dataset

We used lesion volumes and behavioral data for 52 subjects consecutively recruited independent of aphasia diagnosis and tested within an average of four days after unique left hemisphere stroke onset (range 1-12 days; reported in Ding et al., 2020). Lesion maps were processed similarly to the MRRI dataset, see Ding et al. We included two behavioral scores of connected speech: words produced per minute and proportion pronouns to nouns produced during narrative storytelling.

### Predictive Validity Comparison

Implementation of the PVC method involved two phases: model building and model comparison. We illustrated each phase using behavioral data simulated from lesions to distinct Brodmann regions (see Method sub-section *Simulating behavior from lesion volumes* for details on the data generating process). PVC was formulated to take advantage of the software toolkit LESYMAP, which provides access to several different lesion behavior mapping techniques (Pustina et al., 2018). We focused on SCCAN (sparse canonical correlation analysis for neuroimaging) because of its performance on both real and simulated data, its ability to perform well with typical sample sizes, and its robustness to correlated damage across regions (Pustina et al., 2018). SCCAN also generates interpretable (sparse) LBMs alongside explicit predicted values for behaviors and is designed to find the maximal (linear) correlation between the lesion patterns and behavior. We note that other multivariate mapping methods that produce behavioral predictions could also be used with the PVC approach to compare brain damage-behavior associations. To demonstrate PVC’s capacity to successfully use other methods, we used support vector regression (SVR) as a secondary method for analyzing both the MRRI and Schnur Lab datasets.

### Model Building

In the Model Building phase of the Predictive Validity Comparison (PVC) method, the first step (Figure 1A) is to collect segmented lesion volumes for each subject, aligned within a common coordinate system (*e.g.*, MNI space). Next, the two behavioral scores (B_1_, B_2_; Figure 1B) are *z-*scored (mean centered, scaled by the standard deviation), producing the normalized behaviors Z_1_ and Z_2_. Normalization is critical to ensure that simple scale differences between the variables do not influence the lesion-behavior map (LBM) fitting process.

**Figure 1.**
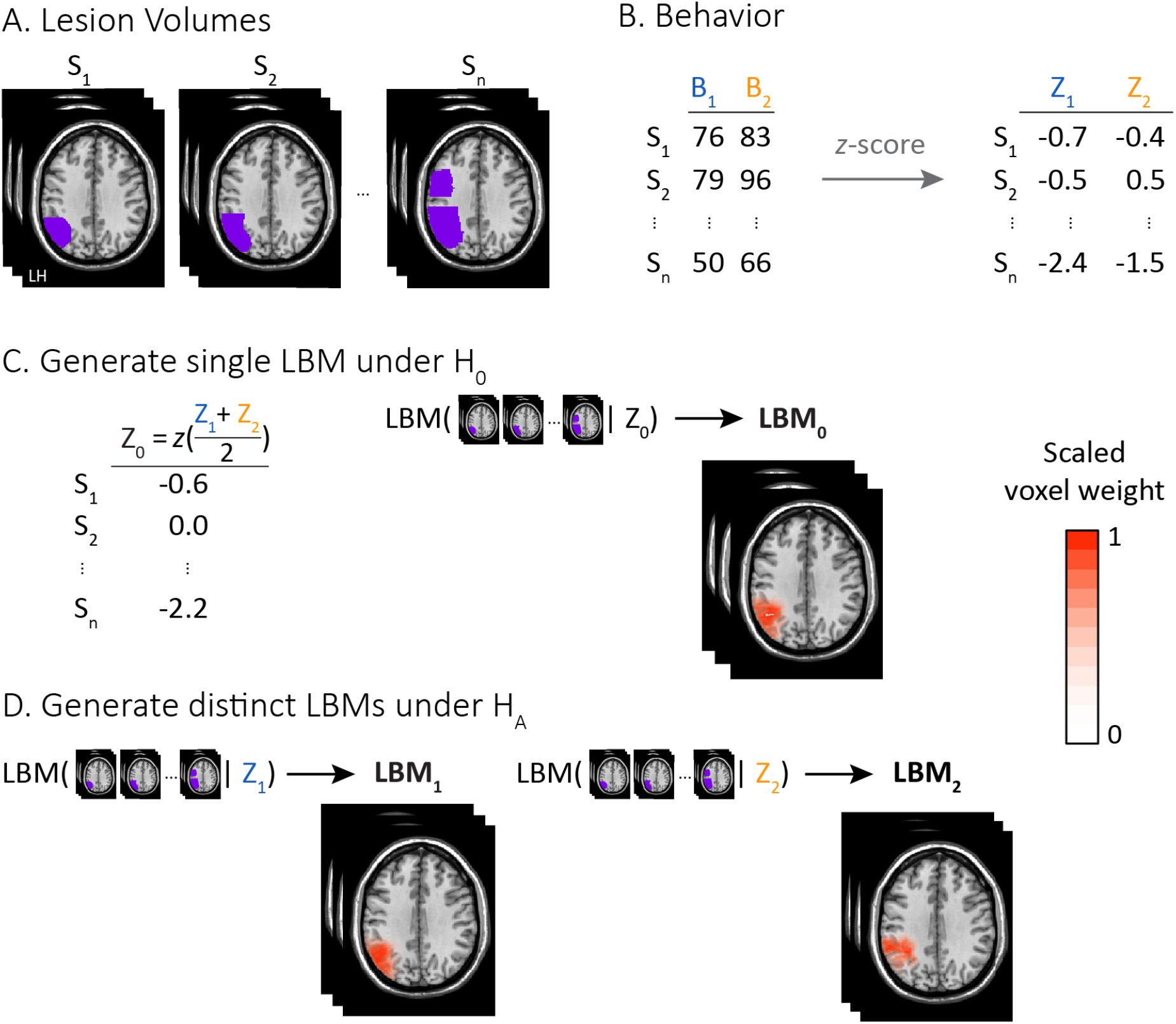
Model Building phase in the Predictive Validity Comparison (PVC) method. **A.** Normalized, segmented lesion volumes for all participants (S^1^ to S_n;_ purple denotes lesioned voxels). **B.** Center and scale (z-scored) behavioral scores (B_1,_ B_2_) prior to fitting for S_1_ to S_n_ (B_1_ → Z_1_, blue; B_2_ → Z_2_, orange). **C.** Under the Null Hypothesis (H_0_), a single LBM is fit using a single behavior score, *Z*_0_ (z-scored average of Z_1_ and Z_2_, left) and subject’s lesion volume (middle), producing a coefficient for each voxel (scaled from 0 to 1 for display purposes, transparent white to opaque red; right). **D**. Under the Alternative Hypothesis (H_A_), distinct LBMs are fit using participants’ lesion volumes paired with each scaled behavior Z_1_ (LBM_1_) and Z_2_ (LBM_2_).

Next, PVC generates multivariate LBMs under the null and alternative hypotheses. In principle, any technique that produces behavioral predictions is amenable for analysis with PVC. Under the null hypothesis, individual differences in normalized behaviors Z_1_ and Z_2_ arise from a single pattern of lesion damage. Therefore, Z_1_ and Z_2_ measure a single underlying factor and (assuming Z_1_ and Z_2_ are equally predictive of this underlying factor) can be averaged into a single behavioral score Z_0_, which is *z*-scored (Figure 1C). Lesion volumes and Z_0_ are then used to generate a multivariate LBM, denoted LBM_0_. In contrast to the null hypothesis, under the alternative hypothesis (Figure 1D), individual differences in behaviors Z_1_ and Z_2_ arise from distinct lesion patterns and thus PVC creates distinct multivariate LBMs: LBM_1_ using the lesion volumes paired with Z_1_ (Figure 1D, left) and LBM_2_, using the lesion volumes paired with Z_2_ (Figure 1D, right). Together, the PVC method generates three LBMs, one based on the null hypothesis, LBM_0_, and two based on the alternative hypothesis LBM_1_ and LBM_2._

### Building null and alternative models with SCCAN

To ensure the predictions from the LBMs are directly comparable, all hyperparameters must be the same for LBM_0_, LBM_1_, and LBM_2_. For the tests reported here, we chose values based on the real data results and recommendations for best performance from Pustina et al. (2018): sparseness = −0.3 (30% of voxels allowed non-zero beta values; voxel-weights are directional); robust = 0 (do not rank transform input); sparse decomposition iterations = 15. Voxel-weight maps are typically thresholded such that values with less than 10% of the maximum fitted weight are not shown. However, all non-zero voxels are still used in the generation of the predicted behavior scores during model comparison (see *Model Comparison*). The PVC method uses a single (user-controllable) level for sparseness rather than attempting the computationally intensive task of jointly optimizing sparseness across the three datasets. We show (Results sub-section *Sensitivity to hyperparameter choice*) that this choice does not impact the results from the PVC method.

### Building null and alternative models with SVR

As with the SCCAN models, the null and alternative models are fit with the same hyperparameters for support vector regression (SVR). We used the default SVR parameters for LESYMAP (cost = 1, epsilon = 0.1), except we used a linear kernel (rather than a radial basis function) to further reduce the likelihood of overfitting the data.

### Model Comparison

In the Model Comparison phase (Figure 2), we generate predicted behavior scores using the LBMs from the Model Building phase under both the null (LBM_0_) and alternative hypotheses (LBM_1_ and LBM_2_). For each subject, we take their vectorized lesion volume (Figure 2A) and use the fitted LBMs to make predictions for each behavior. The exact calculations for generating predicted values will depend on the method used for generating the LBMs. For LBMs fitted with SCCAN, the across-subject 0/1 lesion matrix (subjects as rows, voxels as columns) is multiplied by the fitted LBM (a column vector of weights). These predictions are then *z*-scored to convert the predictions to a common scale with the behavioral input. Under the null hypothesis, there is only a single fitted LBM (LBM_0_; Figure 2B), and therefore the predicted values for a given subject’s normalized behaviors (Z_1_ and Z_2_) are the same (Figure 2C). Total predictive accuracy is summarized using the total Akaike Information Criterion (AIC) across all subjects and both behaviors. Under the alternative hypothesis, the distinct LBMs (Figure 2D) produce distinct predictions across Z_1_ and Z_2_ for each subject (Figure 2E). AIC is used to measure model fit.

**Figure 2.**
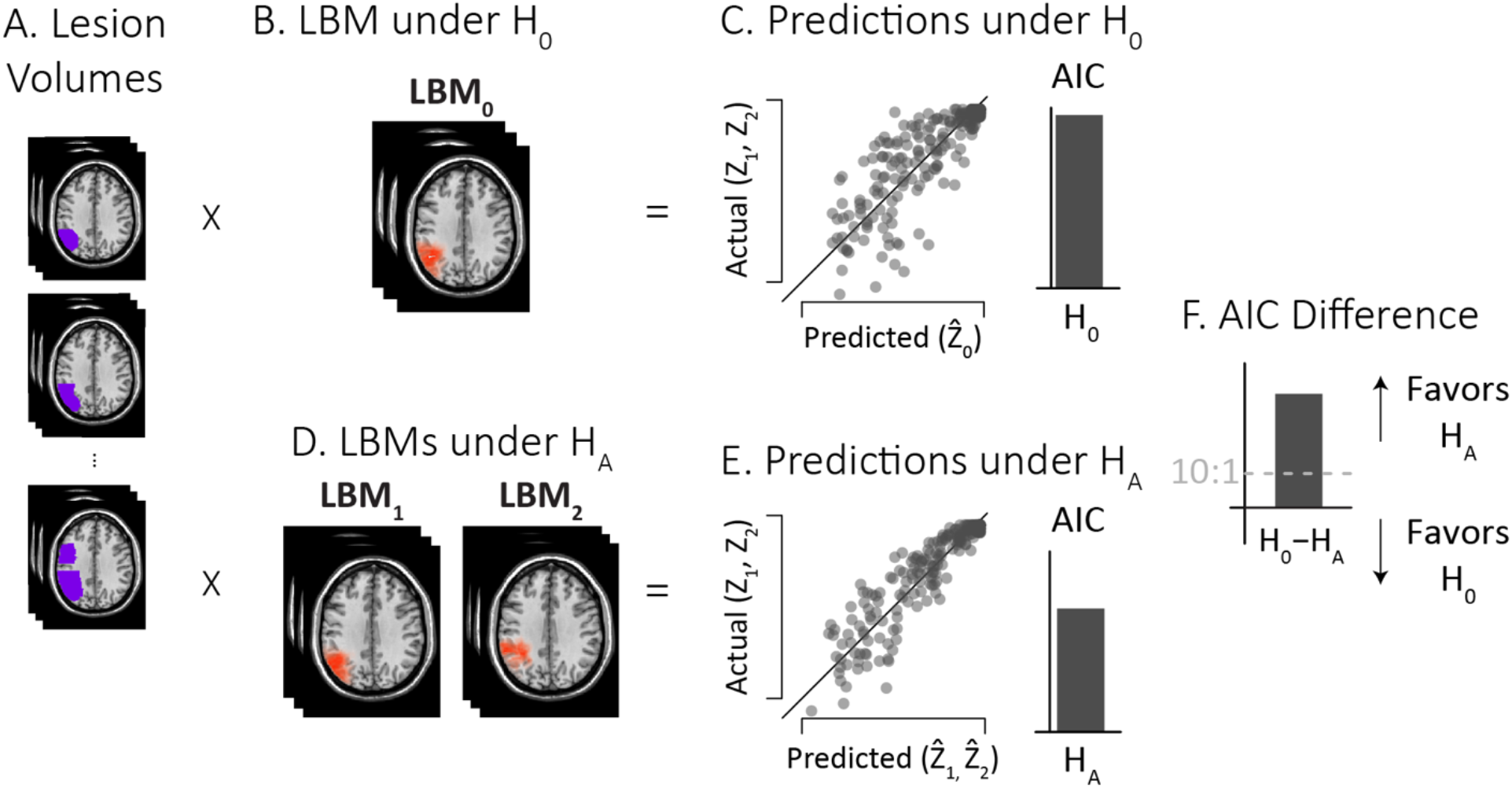
Model Comparison in the Predictive Validity Comparison (PVC) method. **A**. Participants’ lesion volumes are multiplied by each fitted lesion-behavior map (LBM) to generate predicted values. **B.** Under the null hypothesis (H_0_) there is a single LBM, producing a single prediction (Z^^^_0_) for the two actual scaled behaviors (Z_1_, Z_2_). **C**. Predictions are compared with the actual scaled behaviors using AIC (solid line indicates equality between predicted and actual values). The total AIC (inset solid bar) for H_0_ calculated. **D**. Under the alternative hypothesis (H_A_), there are distinct LBMs (LBM_1_, LBM_2_), producing distinct predictions (Z^^^_1_, Z^^^_2_) for the two actual scaled behaviors. **E**. Predictions are compared with actual behaviors and total AIC for H_A_ calculated (inset). **G**. AIC difference (H_0_ – H_A_) determines the winning hypothesis: positive values favor H_A_; negative values favor H_0_. The gray dashed line at +10:1 is the cutoff for claiming decisive support for H_A_.

Because support vector regression (SVR) models are able to fit their training data to an arbitrarily high degree, we used 10-fold cross validation to generate behavioral predictions under the null and alternative hypotheses. AIC was used to measure the overall fit between the null and alternative models.

To compare AIC between the null and alternative hypotheses (Figure 2F), we used the popular BIC difference thresholds (Kass & Raftery, 1995): differences greater than 10 are *decisive* evidence for the alternative hypothesis (behaviors arise from distinct lesion patterns); differences between 6 and 10 are taken as s*trong* evidence for the alternative hypothesis. Negative AIC differences suggest evidence for the null hypothesis (behaviors arise from a single lesion pattern) and the same cutoffs (now negative differences rather than positive) are used. AIC difference values between –6 and +6 may be considered *inconclusive* rather than favoring the null hypothesis because the alternative model has already been penalized for its extra free parameters (see *Calculating AIC*).

### Calculating AIC

AIC measures the total model fit error by summing prediction error across participants for each behavior. Calculating AIC requires choosing a (log) likelihood function (what is the probability of a set of predicted behavioral scores, given the true behaviors) and a value for the penalty term, *k* (how much to penalize extra parameters in the model). For the likelihood function, we assumed the normalized behavior scores followed a normal distribution centered on the true behavior with standard deviation, σ = 0.23. We chose this standard deviation value based on the measured standard deviation when simulating a single behavioral score. We calculated the log-likelihood of the predicted values given the true behavior in R using the *dnorm* function.

Second, to determine the penalty term (*k* in the AIC formula), we used the number of uniquely predicted values. Under the null hypothesis, only a single value is predicted for each subject, therefore *k* is set to N (where N is the sample size); for the alternative hypothesis the penalty term is 2 * N because each behavior score was predicted twice. We chose *k* to be based on the number of predicted subjects rather than the number of fitted voxels because the number of fitted voxels may be affected by choice of hyperparameters, even if predictive accuracy is unchanged (Pustina et al., 2018). Additionally, choosing *k* based on the number of subjects allows us to calculate the relative degrees of freedom between the null and alternative LBMs independently of the method chosen for fitting them. This independence allows us to bypass computationally expensive model optimization when the number of LBMs required to fit the data is yet to be determined. Future work may identify better values for the standard deviation or the penalty term. However, the success of PVC using both SCCAN and SVR (see Results) suggests that the chosen values are generally useful.

### Controlling for lesion volume effects

Prior to scaling the behaviors and fitting the LBMs, the PVC app allows the user to normalize the data based on individual differences in lesion volume. Following Pustina et al.’s (2018) recommendation for multivariate LBM methods, we did not apply lesion-volume correction. The PVC app provides support for lesion-volume correction (regressing out lesion volume from the behaviors) as well as total direct lesion volume control (scaling the lesion matrix by the square root of total lesion volume; Zhang et al., 2014). As noted by Thye et al. (2021), regressing out lesion volume from behavior is often too aggressive (voxels incorrectly excluded from the LBM), because the global lesion volume can be correlated with the lesion volume in the area of interest, leading to missed critical regions (*cf*. DeMarco & Turkeltaub, 2018; Sperber, 2022).

### Method Assessment

We first compared the accuracy of the Predictive Validity Comparison (PVC) method against the two most popular lesion-behavior map (LBM) comparison methods, the Overlap method and the Correlation method, using real data and a range of hyperparameter values. Next, we conducted extensive simulations to assess the accuracy of each method using simulated behavior with a known ground truth (behaviors generated from a single *vs*. distinct lesion patterns).

### Assessing the PVC method with real data

We compared the output of the PVC method with the Overlap method and the Correlation method using both the MRRI and Schnur Lab datasets. Each dataset includes a normalized lesion volume per participant and two associated behaviors for each participant. As a first step, we measured the Pearson correlation between the two behavioral scores to indicate the *a priori* plausibility of being able to distinguish between models predicting each behavior—behaviors that are perfectly correlated should be explainable by a single LBM; uncorrelated behaviors should not be explainable by a single LBM. Next, we used SCCAN to fit multivariate LBMs under the null hypothesis (behaviors arise from a single lesion pattern; LBM_0_) and alternative hypothesis (behaviors arise from distinct lesion patterns; LBM_1_ and LBM_2_). For the Overlap method as is commonly done, we visualized slices of maximal distinction between LBM_1_ and LBM_2_ but also quantified the degree of similarity with the Dice coefficient, 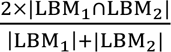: twice the count of non-zero voxels in the intersection of LBM_1_ and LBM_2_, divided by the sum of the count of non-zero voxels in each LBM (0.0 indicates no overlap; 1.0 indicates perfect overlap). For the Correlation method, we visualized the non-zero voxel weights for LBM_1_ and LBM_2_ using a scatterplot and quantified their relationship using the Pearson correlation. For the PVC method, we visualized the predicted vs. actual behavior under the null and alternative hypotheses and used AIC difference to select the best-fitting model.

One problem with the voxel-wise correlation method is the presence of spatial autocorrelation which biases the results. To ensure we tested the PVC method against a statistically-robust alternative, we also estimated an ROI-level correlation measure. For this measure, we averaged the voxel weights in each Brodmann area (using *labelStats* in ANTsR) and then estimated the correlation across the 42 regions. Only regions with non-zero mean weight in at least one map were included in the correlation. By averaging nearby voxels, the ROI-level correlation reduces the effect of the spatial autocorrelation and lowers the degrees of freedom for the statistical hypothesis test.

There are two problems with the Dice coefficient (cf. Ivanova et al., 2021). First, it requires binarizing the data, which destroys information about the relative magnitude of the voxel weights. One alternative to the Dice coefficient that does not require binarizing the data is the one-sided Kuiper test used in Ivanova et al. (2021), but this approach requires having a known “true” brain-behavior locus, rendering it not applicable for general use. However, by including the Correlation method, we do provide a comparator for the PVC that considers the voxel weights directly. The second problem with the Dice coefficient is that it may be affected by the number of voxels in the fitted lesion maps. Using real data, we explicitly manipulated voxel count by varying the sparsity of the fitted maps and assessed the impact on the Dice coefficient.

### Assessing the methods with simulated data

We conducted our simulations using behavioral data generated from a real lesion distribution (*n* = 131 lesion volumes from the MRRI dataset). For each simulation, we selected two (potentially identical) Brodmann regions and generated a behavioral score for each participant based on the lesion proportion within each region. We used the Brodmann atlas provided by LESYMAP (Pustina et al., 2018) for determination of which voxels in MNI space mapped to which Brodmann region. All possible pairs of Brodmann regions in the atlas (41 regions) were included in the simulations, resulting in 820 different-region simulations (the alternative hypothesis was true) and 41 same-region simulations (the null hypothesis was true). Behavioral scores and lesion volumes were then input to the PVC method for analysis. For the Overlap and Correlation methods, we used the LBM_1_ and LBM_2_ output generated as part of fitting PVC.

Kimberg et al., (2007) note that both within- and between-region lesion distributions affect the statistical power of lesion-based analyses. First, it is impossible to draw conclusions about the impact of a brain region on behavior if the is damaged in only one or two subjects; therefore, we excluded simulation runs when both regions had less than 5% patients with at least 5% damage (*n* = 66 simulation runs were dropped, leaving 765 different-region simulations and 30 same-region simulations). Second, because lesion damage covaries across regions (and patients), regions that have identical (un)damage cannot be independently assessed for their effect on the studied behaviors. Therefore, we explicitly examined how the between-region damage correlation impacted the validity of each method.

For the example data shown in Figures 1 and 2, we used the simulation results from Brodmann regions 39 and 40. These regions were chosen because they had adequate lesion loads (more than 45% of patients had at least 5% lesion load in each region) and a moderate-to high between-region lesion-load correlation (*r* = 0.64) which reflects real-world usage and non-trivial lesion-overlap.

To compare the results of each method using the simulated data, we first chose thresholds corresponding to decisions in favor of H_0_ (maps are the same) *vs*. H_A_ (maps are different). For PVC, we used the AIC difference: greater than +10 supports H_A_, less than −10 supports H_0_, inconclusive otherwise. For the voxel-wise and ROI-level correlations: greater than 0 and *p* < 0.05 supports H_0_; otherwise supports H_A_. We chose to use statistical significance rather than a particular value for *r* for simplicity and because previous work employing the method does not provide a clear threshold for “large” correlations when considering LBM similarity (*e.g.*, Zhang *et al*., 2014; Wiesen *et al*., 2019). For the Overlap method: Dice coefficient less than 0.5 supports H_A_, greater than 0.5 supports H_0_. Although there is not a statistically justified, commonly used threshold for the Dice coefficient, 0.5 had intuitive appeal as it judges maps as to whether they share the majority (> 0.5) of their voxels or not (< 0.5). We discuss the problems engendered by a lack of a statistical threshold for the Dice Coefficient in the General Discussion.

### Simulating behavior from lesion volumes

To generate the behaviors, each simulation assumed behavior scores were linearly related to the proportion of damaged voxels in the region assumed to be responsible for that behavior (100% damage corresponds to 0% accuracy). We refer to this damage proportion as the participant’s lesion load for that region. Second, we assumed scores were probabilistic because of binomial-type noise, that is, the variation in behavior was proportional to the uncertainty of an incorrect *vs*. correct response. We adopted this approach because it is more reflective of human choice behavior. Individuals with an expected accuracy near 50% show more variability than individuals with an expected accuracy near 0% or 100%. This assumption was equivalent to assuming that individuals have a “true” behavioral score (say 75% correct), but varied from this number because of finite sampling, just as a few tosses of a fair coin would not be expected to produce an exactly equivalent number of heads and tails. See Supplemental Figure S1 for a graphical overview of this process.

Generating behavioral scores for each participant therefore required two steps (repeated for each behavior). First, the true behavior score was set to their lesion load (*L*, from 0-1) in the selected region. Next, we added noise to this value proportional to L*1-L. This procedure produces more noise for lesion loads close to .5 and less noise for lesion loads close to 0 or 1. For subjects with 0% or 100% damage to a region, we added a minimal amount of noise to the behavior (between *+* and - 0.025). Variability in behavior was necessary as no variability would make determination of “same” LBMs trivial (the behaviors and thus the fitted LBMs would be identical). Simulated behaviors were truncated to be within 0 and 1 for all participants.

### Output provided by the PVC App

As part of developing the PVC method, we built an R Shiny Application that implements it (https://sites.google.com/site/ttschnur/researchprojects/predictive-validity-comparison-for-lesion-behavior-mapping). This application provides several outputs. First, the behavioral data are output before and after normalization (including lesion-volume-correction if requested) along with the predicted behavioral data in CSV files. The SCCAN-fitted LBMs for behavioral scores B_0_, B_1_, and B_2_ are output as compressed NIFTI files (.nii.gz). The average lesion volume (proportion of patients with damage at each voxel) and the lesion *mask* volume (thresholded average lesion volume) are also output as compressed NIFTI files. Finally, the input parameters and numeric output are included in a human- and machine-readable YAML file: behavioral variable names, subject identifier, mask threshold, AIC values, SCCAN iteration number, and SCCAN sparseness value. SCCAN voxel directionality is encoded by the sign of the sparseness value where negative indicates “directional” weights (Pustina et al., 2018).

## Results

We developed a novel method for determining whether two behaviors are better predicted by a single lesion-behavior map (LBM) *vs*. distinct maps. Because of its emphasis on the predictive power of LBMs, we termed the method Predictive Validity Comparison (PVC). We first used two real data sets to assess the practical utility of the PVC method against the two most common methods for LBM comparison: the Overlap method and the Correlation method. Next, we used simulated data to determine the sensitivity (correctly detecting when behaviors are generated from distinct lesion patterns) and specificity (correctly detecting when behaviors are generated from a single lesion pattern) of the three methods across a range of hyperparameters.

### Assessing the PVC method with real data

### MRRI data

We applied the PVC, Overlap, and Correlation methods to the MRRI data (see Figure 3), which comprised responses from two behaviors, the Western Aphasia Battery Aphasia Quotient (WAB-AQ) and accuracy on the Philadelphia Naming Test (PNT) collected from 130 participants with chronic left-hemisphere stroke. First, we compared the behaviors to determine the *a priori* plausibility of using a single vs. distinct LBMs for explaining the individual variation in each behavior. As shown in Figure 3A, the behaviors were extremely highly correlated across subjects (*r* = 0.89, *p* = 10^-16^). Under the null hypothesis (H_0_) that individual differences across behaviors arise from a single lesion pattern, we fit a single LBM (LBM_0_) to the averaged behaviors (Figure 3B, *left*). Under the alternative hypothesis (H_A_) that individual differences arise from distinct lesion patterns, we fit two LBMs (LBM_1_ and LBM_2_), one for each behavior (Figure 3B, *right*).

**Figure 3.**
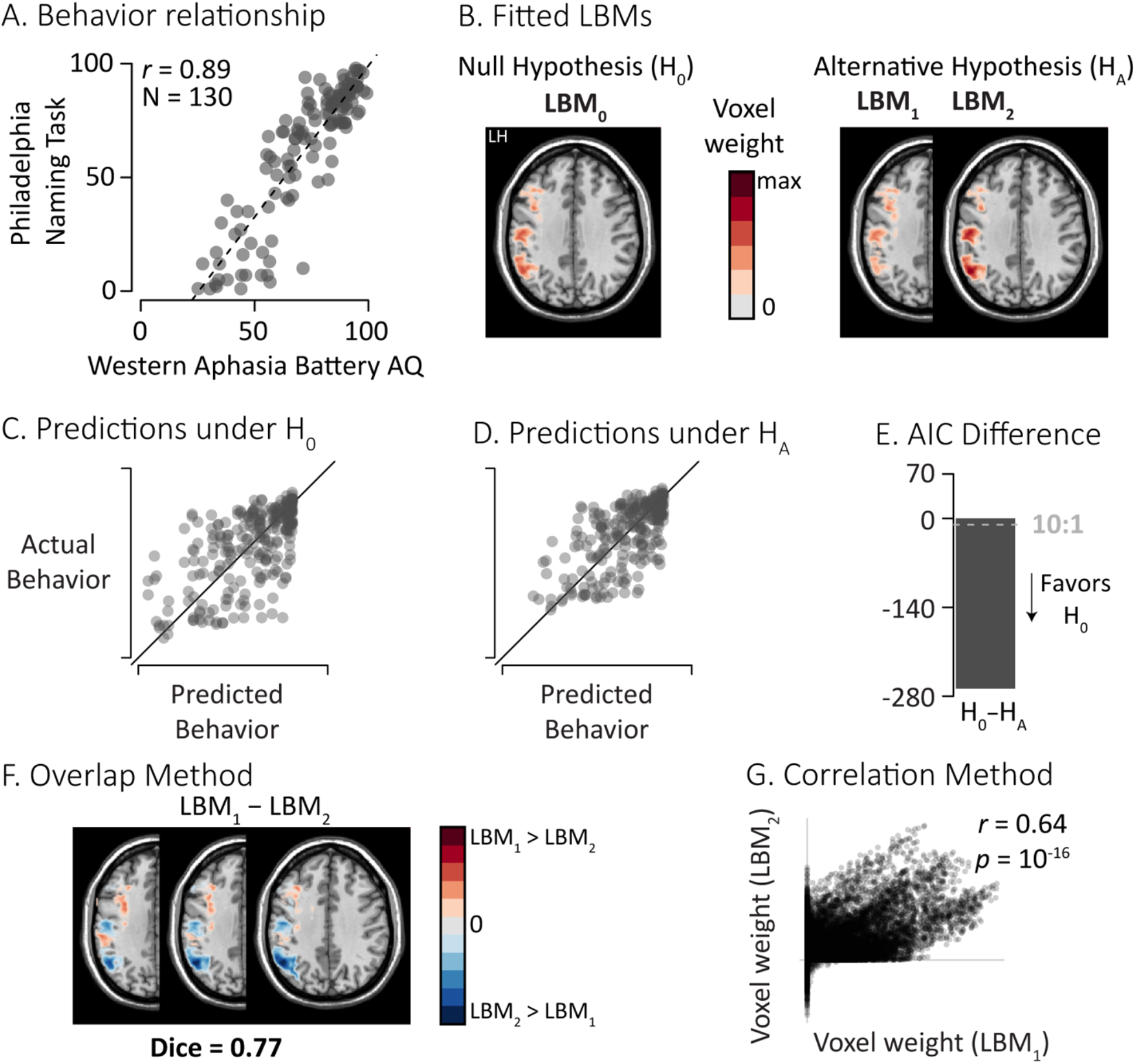
Comparing lesion-behavior maps (LBMs) built from the MRRI data. **A.** Scatter plot showing the strong linear relationship (*r* = 0.89) between the Western Aphasia Battery Aphasia Quotient (AQ) and the Philadelphia Naming Task (% accuracy) for 130 participants. **B**. Fitted voxel weights (max-scaled across maps; transparent white to opaque red) from LBMs fitted under the null hypothesis (LBM_0_) and alternative hypothesis (LBM_1_ and LBM_2_). The axial slice shown (z = 107) had the most non-zero voxel weights for both LBM_1_ and LBM_2_. LH: Left Hemisphere. **C./D.** The PVC method compares the actual behavior to predictions generated under the null hypothesis (H_0_) and the alternative hypotheses (H_A_). Solid diagonal line indicates perfect prediction. **E.** The AIC difference was decisive for H_0_ (cutoff at −10, gray dashed line). **F**. The Overlap method highlights slices that differ between the LBMs. Voxel color indicates the sign and magnitude of the difference (opaque blue to transparent white to opaque red; max-scaled). The dice coefficient of 0.77 suggests a moderate to high overlap between the maps. **G**. The Correlation method suggested a moderate relationship (*r* = 0.64) between the weights in LBM_1_ and LBM_2_.

The PVC method explicitly compares the LBMs generated from null and alternative hypotheses. Under the null hypothesis (H_0_), a single LBM was fit across behaviors and used to produce a single predicted value for both the *z*-scored WAB AQ and PNT scores for each participant. Despite this constraint, predicted values were a reasonable match for the actual values across participants (R^2^ = 0.42; Figure 3C). Under the alternative hypothesis (H_A_), distinct LBMs were fit to each behavior and distinct values were predicted for each participant’s WAB AQ and PNT scores; overall accuracy under H_A_ was also reasonable (R^2^ = 0.45; Figure 3D). We used AIC to statistically compare the predictive accuracy of H_0_ and H_A_, finding decisive support for H_0_: individual differences across WAB AQ and PNT are best explained by a *single* LBM (AIC Difference = −268, in favor of H_0_; Figure 3E). Using multivariate LBMs generated with SVR rather than SCCAN produced qualitatively similar results, *i.e.*, decisive support for H_0_ (see Supplemental Figure S2 A-C).

For the Overlap method, we first visualized the difference map (LBM_1_ – LBM_2_) to inspect the spatial non-overlap between the fitted LBMs. Although the fitted LBMs showed some similarity upon visual inspection (*cf*. Figure 3B, right) there were also areas of non-overlap, resulting in a moderate to large Dice coefficient of 0.77 (Figure 3F).

For the Correlation method (Figure 3G), the correlation between the voxel weights for LBM_1_ and LBM_2_ was moderate, positive, and statistically significant (*r* = 0.64, *p* = 10^-16^). Correcting for spatial autocorrelation using the ROI-level correlation (*df =* 40 instead of *df =* 233,567 for the voxel-level correlation) yielded a smaller but still significant correlation between the maps (*r* = 0.56, *p* = 0.0005).

### Schnur Laboratory data

Next, we compared the three comparison methods using two unrelated behaviors calculated from narrative storytelling (words produced per minute and proportion pronouns to nouns) collected from 52 subjects during the acute stage of left hemisphere stroke (Ding et al., 2020). Unlike the MRRI data, the two behaviors in the Schnur Lab data had little to no linear relationship (Figure 4A; *r* = −0.17, *p* = 0.23). Under the null hypothesis that individual differences arose from a single lesion pattern, we fit a single LBM across behaviors (Figure 4B); for the alternative hypothesis, distinct LBMs were fitted to each behavior (Figure 4B, right). For the PVC method, the predictions under each hypothesis were noticeably different. Under the null hypothesis (H_0_), the accuracy of the predictions was poor (Figure 4C, R^2^ = 0.12), whereas for the alternative hypothesis (H_A_), the accuracy was better (Figure 4D; R^2^ = 0.48). Even after accounting for the flexibility of fitting distinct LBMs, the PVC method decisively favored the alternative hypothesis (Figure 4E; AIC difference = 1143, in favor of H_A_). Using multivariate LBMs generated with SVR rather than SCCAN produced qualitatively similar results, *i.e.*, decisive support for H_A_ (see Supplemental Figure S2 D-F).

**Figure 4.**
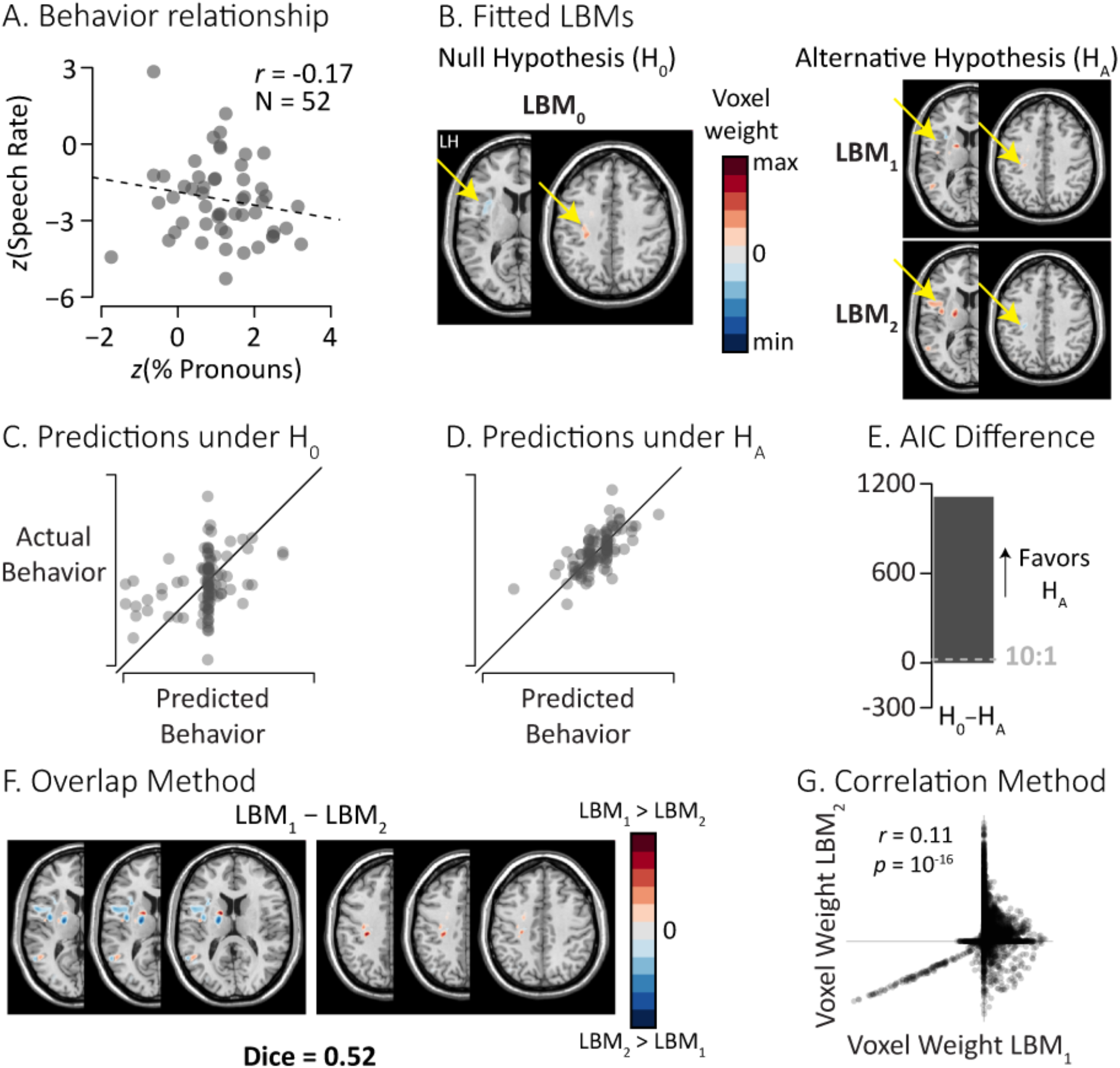
Comparing lesion-behavior maps (LBMs) built from the Schnur Laboratory data. **A.** Scatter plot showing no linear relationship between z-scored words per min (Speech Rate) and z-scored relative percentage of pronouns to nouns produced for 52 participants. **B**. Fitted voxel weights (max-scaled across maps; transparent white to opaque red) from LBMs fitted under the null hypothesis (LBM_0_) and alternative hypothesis (LBM_1_ and LBM_2_). The axial slices shown (z = 84, 113) had the most non-zero voxel weights for LBM_1_ and LBM_2_, respectively. LH: Left Hemisphere. Yellow arrows note areas of distinction across the slices. **C./D.** The PVC method compares the actual behavior to predictions generated under the null hypothesis (H_0_) and the alternative hypotheses (H_A_). Solid diagonal line indicates perfect prediction. **E.** The AIC difference (H_0_ – H_A_ = 1143) used by the PVC method was decisive for H_A_ (cutoff at +10, gray dashed line). **F**. The Overlap method highlights slices that differ between the LBMs. Voxel color indicates the sign and magnitude of the difference (opaque blue to transparent white to opaque red; max-scaled). The Dice coefficient of 0.52 suggests a moderate overlap between the maps. **G**. The Correlation method assesses the relationship between the weights in LBM_1_ and LBM_2_, suggesting a weak but significant positive correlation (*r* = 0.11, *p* = 10^-16^).

In contrast to the clear results from the PVC method, both the Overlap method and the Correlation method produced more ambiguous results. For the Overlap method (Figure 4F), the difference map showed some distinct regions of non-overlap, but there was still a moderate degree of overlap (Dice coefficient = 0.52). For the Correlation method, despite that lack of a behavioral correlation, there was still a small positive correlation in the voxel weights (*r =* 0.11, *p* = 10^-16^). The ROI-level correlation showed a *stronger* positive correlation (*r* = 0.53, *p =* 0.017) despite the drastically reduced sample degrees of freedom (*df* = 18 *vs. df* = 38,672).

### Sensitivity to hyperparameter choice

One source of the flexibility in multivariate LBM fitting methods is the presence of tunable hyperparameters. For instance, SCCAN (Pustina et al., 2018) allows users to set (or optimize) a desired level of sparseness for their data and choose whether to allow voxel weights to have directionality (directional weights allow for positive and negative relationships between lesions and behavior across voxels; non-directional weights assume a fixed direction of relationship across all voxels). Because these hyperparameters affect the fitted weights, they will necessarily affect LBM comparison methods derived from these weights. Therefore, we re-analyzed each of the real data sets using a variety of sparseness levels (in SCCAN, higher levels of the sparseness parameter correspond to *more* voxels included in the fitted LBM) while also varying voxel weight directionality (with and without directionality).

Figure 5 shows the effect of sparseness across methods for both the MRRI data set and the Schnur Laboratory data set. For PVC, changing sparseness and voxel directionality had no qualitative impact on the AIC differences; all tested parameter values showed decisive support for the same hypothesis for both the MRRI data (supporting H_0_: the LBMs are the same; Figure 5A) and the Schnur Laboratory data (supporting H_A_: the LBMs are different; Figure 5B). In contrast, the Dice coefficient (Overlap method), voxel-wise correlation, and ROI-level correlation showed strong variation across changes in sparseness and voxel directionality.

**Figure 5.**
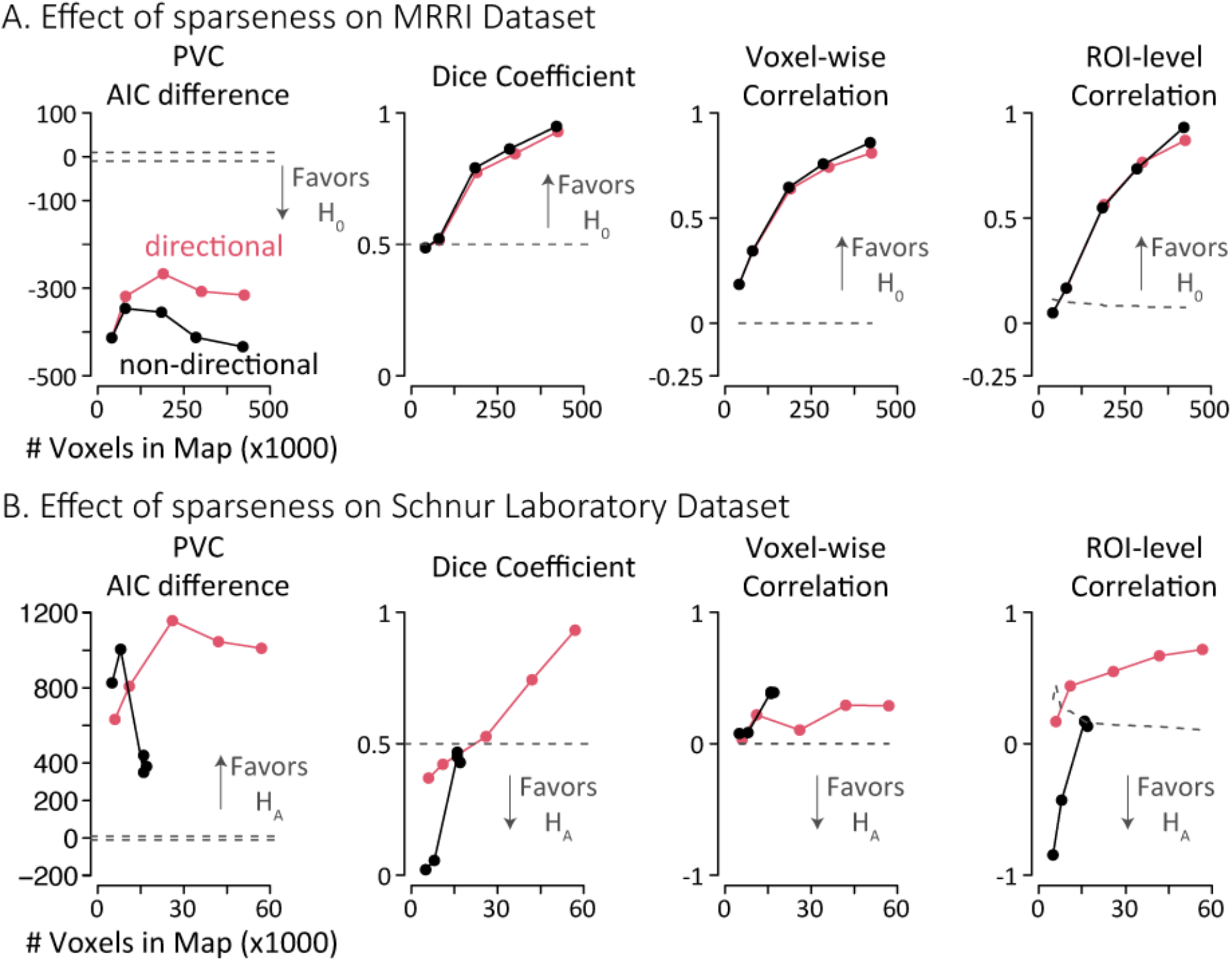
Effect of varying SCCAN sparseness parameter and directionality of voxel weights on lesion-behavior map (LBM) comparison results using real data. More sparse models have fewer voxels in the resulting LBM; number of non-zero voxels in the fitted LBMs are plotted on the horizontal axis (sparsity) and voxel-weight directionality is shown in separate lines (pink: directional weights allowed; black: non-negative weights only). The dashed gray line provides the threshold for determining same *vs*. different LBMs. **A.** For the MRRI dataset (H_0_), changing the sparsity of the result had quantitative impacts on the AIC difference used by the predictive validity comparison (PVC) method (Left), but no change in the conclusion (all simulations decisively supported the null hypothesis). For the Dice coefficient used by the Overlap method (2^nd^ column) and the correlation-based methods (3^rd^ and 4^th^ columns), the sparseness of the model modulated the quantitative metrics, but there was little impact of voxel-weight directionality on the results. Across much of the range, all methods supported the same conclusion (maps are the same), except for sparse maps using the ROI correlation method. **B.** For the Schnur Lab dataset (H_A_), the PVC method again showed quantitative differences across parameter manipulations, but no decisional change (all simulations favored the alternative hypothesis). The Dice coefficient and the ROI-level correlation were inconsistent and strongly dependent on the choice of sparseness and voxel directionality; the voxel-wise correlation showed consistent support for H_0_.

For both datasets, the Dice Coefficient switched from support for the separate-map solution to the single-map solution as the number of voxels in the map was increased. For the voxel-wise correlation, there was a substantial change in the strength of between-map correlation, but relatively consistent support for the single-map solution despite the clear differences between the MRRI and Schnur Laboratory datasets. For the ROI-level correlation, the reduced degrees of freedom led to some inconclusive results for the smallest LBM sizes, but also showed a clear dependency on hyperparameter values, switching from support for the separate-map solution to the single-map solution for the Schnur Laboratory data.

Such strong variation for non-PVC metrics within and between datasets with strikingly different behavioral correlations obviates the ability to a derive a single sufficient cutoff value to determine significant similarity between LBM_1_ and LBM_2_ for the Overlap and Correlation methods, regardless of hyperparameter choice. In contrast, the consistency of the PVC method provides a measure of assurance for PVC users that the specific choice of hyperparameters will not pre-determine the result of the LBM comparison.

### Assessing the methods with simulated data

We assessed the accuracy of each method using behavioral scores simulated from the MRRI lesion data. In each simulation, the ground truth is known: the scores were derived from the amount of lesion to distinct Brodmann regions (ground truth different) or from a single Brodmann region (ground truth same). Because lesion-symptom mapping accuracy varies based on lesion distribution, we binned the different-region simulations by the between-region damage correlation (the correlation between the damage within each region across participants; for ground truth same simulations this correlation is always 1.0).

Figure 6 shows the results for the ground truth “different” simulations. As shown in Figure 6A, PVC performed best overall, showing near-ceiling accuracy (at least 95%) for between-region damage correlations up to around 0.7 before breaking down. Next, the ROI-level correlation performed well at low between-region damage correlations but started to show worsening sensitivity at correlations around 0.25. Similarly, the voxel-level correlation also showed poor sensitivity with increasing between-region damage correlations. It is notable that the voxel-level correlation also showed atypically poor performance for regions with *negative* damage correlations. The Overlap method (Dice coefficient) performed similarly to the voxel-level correlation except that it performed better for negative between-region correlations and worse for high correlations. To highlight the comparisons with PVC, we plotted each model’s performance relative to PVC for conditions in which any method had at least 50% accuracy. Figure 6B shows PVC was better than other methods. Although all models performed relatively poorly for regions whose damage was highly correlated, these occasions were rare. No method can determine separable neural bases if damage between regions is extremely correlated. Lesion distribution is dependent on the specific data set and here was determined by the lesion distribution in the MRRI dataset. Figure 6C shows the relative frequency of the between-region damage correlations. Most regions had moderate to weak damage correlations and high correlations were rare. Thus, for ground truth “different” simulations PVC performed best overall, except when between-region damage correlations were extremely high where all methods failed.

**Figure 6.**
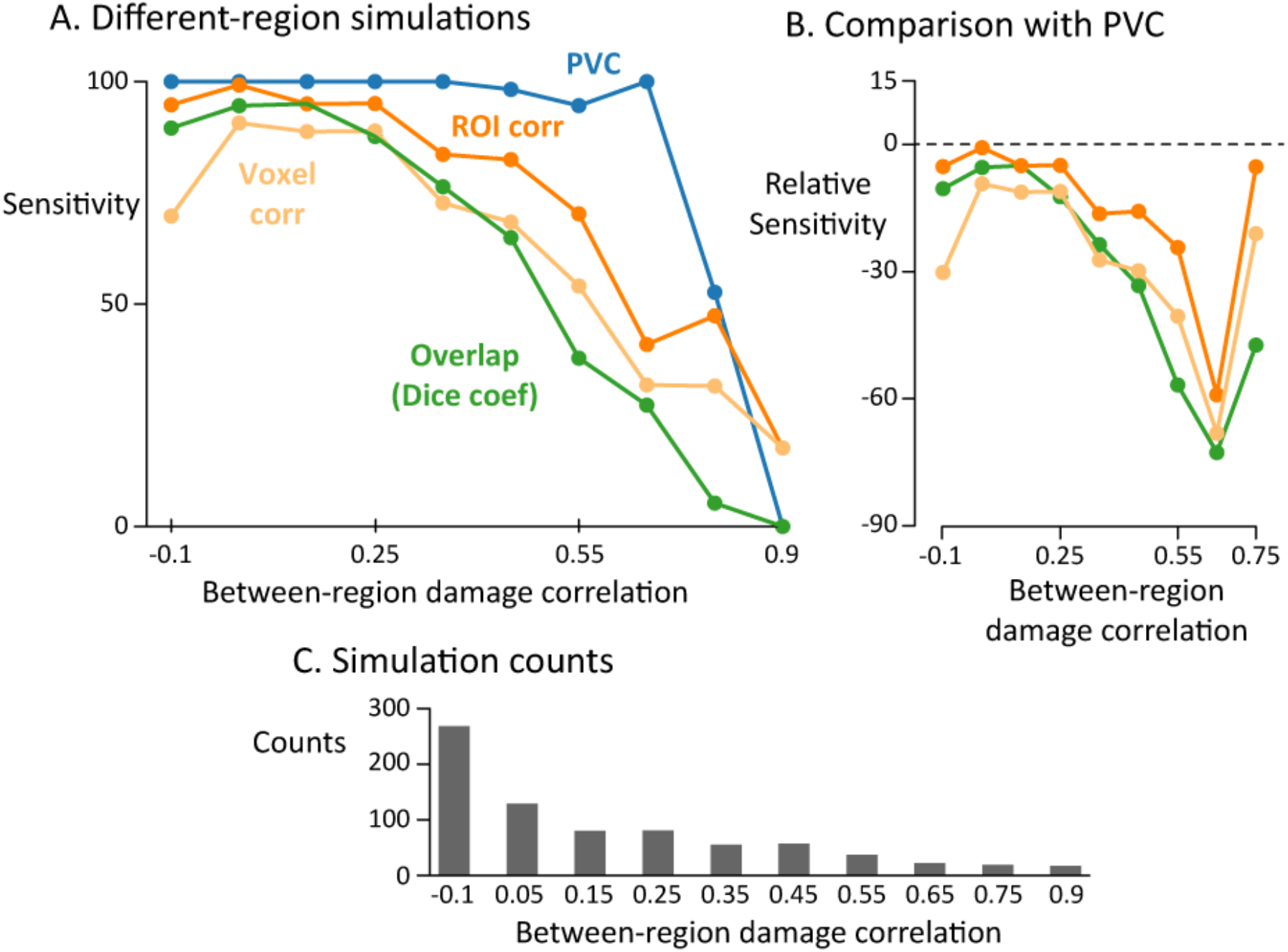
Simulation results. **A.** Averaged accuracy from the different-region simulations provide the sensitivity (% correct for judging that two behaviors have different neural bases) for each method (PVC: Blue; Voxel-wise correlation: pale orange; ROI-level correlation: orange; Overlap method/Dice coefficient: green), binned by the between-region damage correlation. Each point represents the average sensitivity across a varying number of simulations. **B.** Relative sensitivity vs. PVC for each method (same colors as Panel A). Only bins with > 50% sensitivity are displayed. The dashed line at 0 represents the sensitivity of the PVC method (blue line in Panel A). **C.** The number of simulations within each bin (total simulation count = 765). This distribution was determined by the lesion distribution in the MRRI dataset.

The ground truth “same” simulations provide an opportunity to consider the false positive rate of each method: given two behaviors that are subserved by the same neural basis (but may still differ due to within-subject variability), how often does the method claim that distinct maps are needed to accurately predict them? Of the 30 regions that met the minimum-damage inclusion criterion, PVC and the Overlap method were perfect (30/30 correct; 0% false alarm rate) and the Correlation methods nearly so (29/30, 3% false alarm rate). Importantly, although the neural bases for the simulated behaviors are the same, the subsequent fitted maps are not the same because of the noise added to the behavior during the simulation. Thus, using the same-region simulations as a limiting case, we demonstrate that PVC does not always prefer a two-map solution.

Taken together, the results from the ground truth “different” and ground truth “same” simulation show that PVC outperformed all methods across nearly the entire range of between-region damage correlations. All methods suffered poor sensitivity when high between-region damage correlations made distinguishing LBMs impossible. The reason PVC outperforms these methods is because the criterion is directly linked to the performance of the LBM (how well it predicts behavior) rather than surface similarity of the LBMs.

## Discussion

The primary goal of building a multivariate lesion-behavior map (LBM) is to learn the lesion pattern most-predictive of individual differences in a behavior. Often researchers collect two (or more) behaviors from a single cohort and try to determine how LBMs fitted to each behavior are similar or different. A crucial missing step in this process is determining whether multiple LBMs are, in fact, needed to explain individual differences across the behaviors. We developed the novel Predictive Validity Comparison (PVC) method to fill this gap. The model formalizes the task of picking the appropriate number of LBMs to fit by explicitly comparing the predictive power of a single LBM (predictions generated under the null hypothesis) *vs.* distinct LBMs (predictions generated under the alternative hypothesis). Only after the single LBM predictions are found inadequate (rejecting the null hypothesis) are we justified in interpreting the differences in the lesion patterns learned from fitting distinct LBMs to each behavior. To encourage adoption of our novel PVC method for investigating associations between lesion patterns and behavior, we have released an interactive web-based application (along with all source code) at https://sites.google.com/site/ttschnur/researchprojects/predictive-validity-comparison-for-lesion-behavior-mapping.

### Methodological advances

The PVC method offers several advances when applying lesion-behavior-mapping to understand whether different brain regions are necessary for different behaviors. Using two real data sets and extensive simulations, we showed that the PVC method was able to make clear and accurate decisions in favor of single *vs*. distinct LBM solutions. This accuracy and precision contrasted with the less accurate and more variable (i.e., sensitivity to choice of hyperparameters) results from the two most common LBM comparison techniques—the Overlap method (using the Dice coefficient) and the Correlation method (using either voxel-level or ROI-level correlations). Within the PVC method, there is an established method (difference in AIC) for determining whether two behaviors are better explained by a single *vs.* distinct LBMs. For both the Dice coefficient and the voxel-wise correlation (and the ROI-level correlation), there is no single threshold that can accurately determine whether LBMs are the same *vs*. different. We used the statistical significance of the correlation to determine whether two LBMs are distinct (*p* < α, testing for *r* > 0). However, for both of our real datasets, the voxel-wise correlation was significantly greater than 0, even when the correlation between the behaviors was negative. Additionally, the magnitude of the correlation (ignored when thresholding with the *p-*value) would seem to be relevant: *r = 0.9, p < 0.05* is certainly better evidence for the equivalence of two LBMs than *r* = 0.01, *p* < 0.05, but the statistical significance criterion ignores this obvious difference.

Second, the PVC method does not require extensive tuning of hyperparameters (e.g., sparseness, voxel directionality) and is relatively insensitive to their value. In contrast, the Overlap method (Dice coefficient) and Correlation method were strongly dependent on the choice of LBM fitting parameters, obviating the ability to set a single, principled threshold that works across data sets. For instance, changing the sparseness parameter had drastic effects on the Dice coefficient and the LBM correlations, but did not adversely impact the ultimate decision from PVC (single *vs*. distinct LBMs). This difference in hyperparameter sensitivity is caused by the fact that the sparseness parameter has a strong effect on *which* voxels are selected (affecting the Dice coefficient) and on the *size* of the coefficients (small weights are driven to zero, affecting the voxel-wise/ROI-level correlations), but the predictive accuracy is largely unchanged. For LBMs with moderate spatial overlap, reducing the number of voxels in each LBM (more sparsity) can increase the perceived difference between the maps without affecting the accuracy of each LBM. This “feature” of the sparseness parameter is why the authors of SCCAN refer to this parameter as controlling the *interpretability* of the solution rather than its *accuracy* (Pustina et al., 2018). In sum, PVC offers the advantage of less sensitivity to shifting hyperparameters in comparison to other comparison methods.

Third, the PVC method provides excellent specificity and sensitivity in the face of lesion distribution characteristics and several options for accounting for lesion volume. Extensive simulations demonstrated consistently sensitive performance (accurately detecting “different” LBMs) of the PVC method with lesion loads for damage correlations up to around 0.7. Performance on ground-truth “same” comparisons was similarly high. Because PVC is better in comparison to other methods at a wide range of between-region damage correlations, it is more advantageous to use PVC to estimate whether two LSMs are the same or different. In most cases one cannot know in advance which regions will be critical for behavior in order to *a priori* pick a methodological approach based on between-region damage correlation.

### Advancing theories of brain-behavior associations with PVC

The PVC method provides two specific advantages for testing theories of brain-behavior relationships in comparison to other methods. First, PVC provides a principled method for determining whether a single *vs*. distinct LBMs are needed to explain two behaviors by defining a clear, statistical comparison between these two alternatives: whether the data are fit better by a single *vs*. distinct LBMs. In contrast, the popular Overlap and Correlation methods operate by first assuming what they are trying to prove—that each behavior should be fit by distinct maps— and then explaining observed differences. An analogous error would be trying to enumerate all the possible sources of difference between two clinical groups, without first establishing that the two groups are, in fact, statistically different. Avoiding this critical step increases the likelihood of Type I error proliferation and encourages *post hoc* reasoning about group differences. More concretely, an overlap map will nearly always show some spatial differences between LBMs fitted to different behaviors that a clever researcher can explain; however, such spatial differences may be orthogonal to the predictive validity of the LBMs and a generalizable cutoff for the Dice coefficient has not been established (Schwartz et al., 2009). As we demonstrated here, the PVC method clearly adjudicates between single *vs.* distinct LBM solutions while the Overlap and Correlation methods are incapable of the same.

We note an additional approach adopted by several laboratories including ours to determine if two brain-behavior relationships are the same is to simply regress out one behavior from another and then use the residuals to fit an LBM (Lukic et al., 2021; Martin et al., 2021; Thye et al., 2021). Here too, this approach fails to construct clear alternatives. The lack of clear alternatives leads to interpretation difficulties when no statistically significant LBMs are produced, and researchers resort to using the Overlap method and interpreting an intersection map. In extreme scenarios, regressing out one variable from another can even induce a spurious relationship between the residuals and brain damage.

A second theoretical advance PVC provides is a more useful map of brain-behavior associations judged to be the same. A typical approach to compare LBMs judged as “same” is to take some combination of them (e.g., intersection, union, average) and interpret the resulting map. However, this approach does not yield a map capable of producing behavioral predictions and does not provide a quantitative comparison between the single-map and distinct-maps solutions. PVC provides the clear decision rule along with a map that is both interpretable and usable for predicting behavior. A related advantage of PVC is that using the combined behaviors to fit a single, sparse multivariate LBM can reduce nuisance variation in the fitted weights caused by behavioral variability.

### Limitations & Future Directions

As with common null hypothesis testing, we should be cautious about “accepting” the null hypothesis. This concern is mitigated somewhat by PVC’s use of AIC for model comparison, rather than a strict reliance on p-values that are calculated under the assumption that the null hypothesis is true (thus rendering it circular to accept the null hypothesis). In formulating the model, we were careful to say “distinct lesion patterns” rather than “distinct *neural bases”* as we recognize that behaviors may indeed have distinct (that is, separate) neural bases, but due to data limitations (poor behavioral measurements, low-quality structural images, correlations in lesion load) are inseparable in a particular data set. For instance, perhaps one of the behaviors under study has no clear relationship with the observed brain lesion pattern, rendering the distinct-maps solution no more accurate than the single-map solution, even though there is no shared neural basis for the behaviors. In this case, inspection of the overall predictive accuracy would provide a clue for correct interpretation. Or, individual differences across two behaviors may be based on damage to two different regions, but if these regions (nearly) always have co-occurring damage in stroke, they will not be statistically separable by any lesion-behavior mapping. One promising approach to solve this problem is to focus on acute stroke cases, which show smaller lesions (Ding et al., 2020) and may provide better opportunities for dissociating behaviors. Of course, new lesion-behavior data may be able to show the inadequacy of the single-LBM solution (or indeed override previous evidence for the distinct-maps solution). Therefore, it is important to test the predictive validity of fitted LBMs with newly collected data to assess their generalizability.

Although the current formulation of PVC is limited to comparing two behaviors, the idea behind PVC extends naturally to the many-behavior case. PVC simply asks the question: can the behaviors collected be adequately predicted by fewer LBMs than the total number of behaviors. For instance, if you are comparing performance across multiple measures of speech production following brain injury, how many distinct LBMs are necessary to explain a large percentage of variance in the data? One future approach will be to use canonical correlations (the technique at the heart of SCCAN (Pustina et al., 2018)) to estimate the number of necessary LBMs to explain a given amount of variation in a set of behaviors.

Recently, several new advances offer a complement to the PVC method by estimating the best brain map (or set of multi-modality maps) to predict a single behavior. The key difference between PVC and these methods is that PVC determines whether two behavioral scores are better predicted by a single vs. separate lesion-behavior maps by taking as input a single lesion map per subject and two behaviors per subject. In contrast, Salvalaggio et al. (2020) developed a method for comparing behavioral predictions from different types of brain maps (e.g., lesion maps, structural disconnection maps, and functional disconnection maps). They used ridge regression to obtain predicted behavioral scores for each subject based on each map, and then compared the accuracy of each map type to determine the best predictor. This method used only a single behavioral score per subject, however, and did not compare predictive validity between sets of brain maps across behaviors, as is the case in the PVC method. In a similar vein, Siddiqi et al. (2021) recently developed a method that also uses multiple map types. Their method took as input multiple types of brain maps (lesion maps, transcranial magnetic stimulation maps, deep-brain stimulation maps) and a single behavioral score. The critical step in the algorithm converted the brain maps to combine them into a “circuit map”. This aggregate map was then used to predict an individual’s score on a single behavior. While other methods estimate the best-fitting brain map for a single behavior, PVC provides a means to adjudicate whether different behaviors arise from damage to different vs. the same brain regions. Working together, these methods now enable researchers to determine whether two lesion maps are necessary to predict two behaviors (PVC) as well as determine which multi-modality brain maps best predict a single behavior (Salvalaggio et al., 2020; Siddiqi et al., 2021).

## Conclusions

When comparing two lesion behavior maps (LBMs), the most common approaches assume the maps are different and then try to explain apparent differences. This approach falters by not explicitly testing whether a single LBM predicts both behaviors as well as distinct LBMs. Researchers are left without clear criteria for deciding if the LBMs are different and without a way to estimate a single brain-behavior relationship for behaviors that may have the same neural basis. We developed the Predictive Validity Comparison (PVC) method to overcome these problems. We used both real and simulated data to validate its sensitivity (correctly detecting distinct-LBM solutions) and specificity (correctly detecting single-LBM solutions). PVC uses a principled statistical criterion to determine if the behaviors are better predicted by a single *vs*. distinct LBMs, and then provides the user with interpretable maps for *either* conclusion. With PVC, researchers can make stronger conclusions about the neural separability of behaviors. We have released an open-source, GUI-driven software toolkit built on LESYMAP (Pustina et al., 2018) and ANTsR (Avants, 2015) that implements the approach (available online at https://sites.google.com/site/ttschnur/researchprojects/predictive-validity-comparison-for-lesion-behavior-mapping).

## Declarations

### Acknowledgments

The authors wish to thank the Moss Rehabilitation Research Institute for generously sharing their extensive lesion behavior data. We thank Junhua Ding for input when creating and verifying the PVC approach. Parts of this work were presented at the Society for Neuroscience (2019) and the Society for the Neurobiology of Language (2021).

### Data & Code Availability

Anonymized data that support the findings of this study are available by request to the authors and Moss Rehabilitation Research Institute. PVC is publicly available at https://sites.google.com/site/ttschnur/researchprojects/predictive-validity-comparison-for-lesion-behavior-mapping.

### CRediT authorship contribution statement

John Magnotti: Conceptualization, Methodology, Formal analysis, Visualization, Software, Writing – original draft.

Jaclyn Patterson: Writing – original draft. Software.

Tatiana T. Schnur: Conceptualization, Methodology, Writing – original draft, Supervision, Funding acquisition.

## Supplemental Information

**Figure S1.**
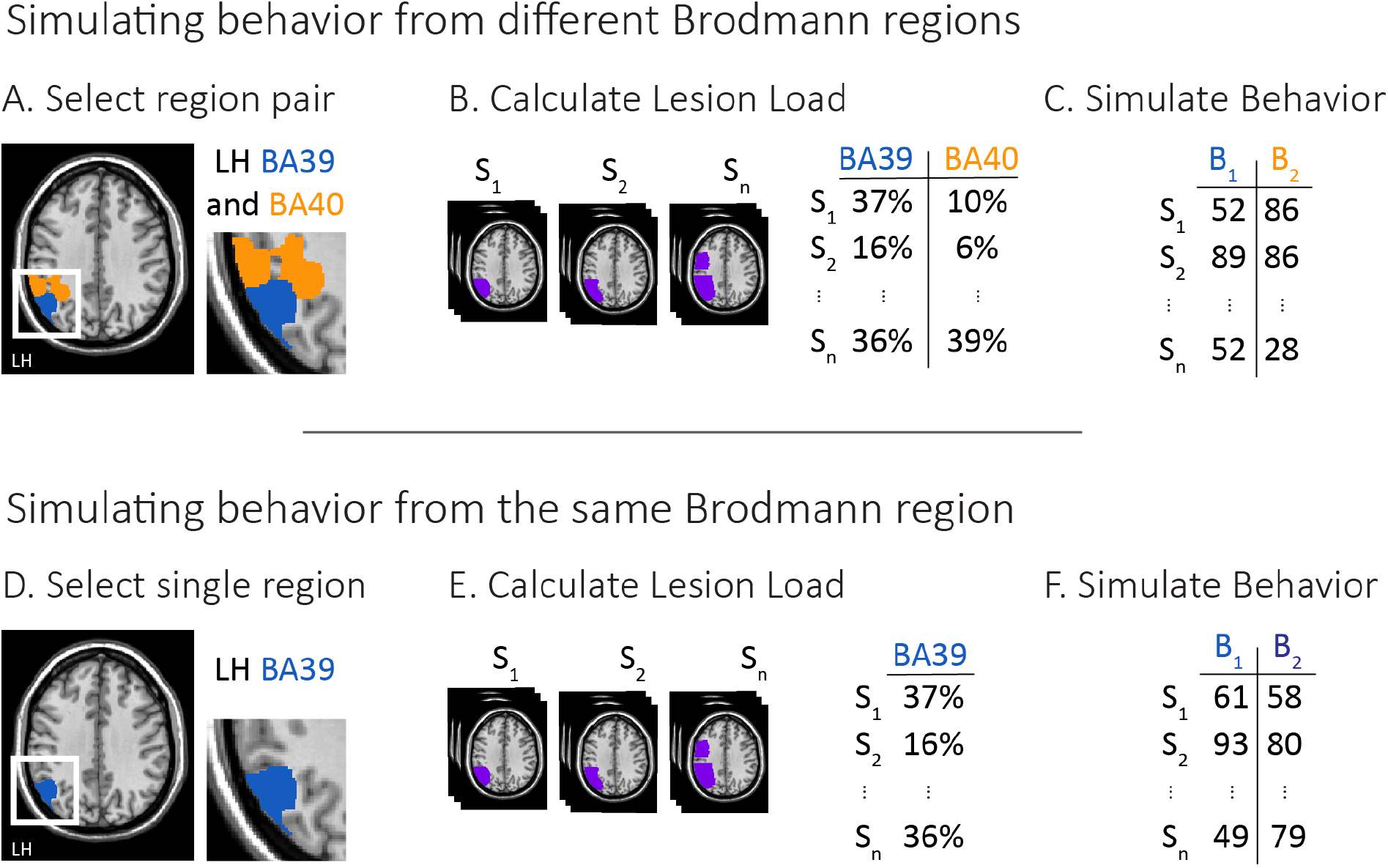
Method for simulating lesion-related behavior using Brodmann regions. **A.** To simulate behaviors B_1_ and B_2_ arising from different Brodmann regions, two region pairs are randomly selected. **B.** Lesion load (L) is calculated for each subject as the proportion of damaged voxels within each region (shown as percentages for display purposes). **C.** Simulated behavior is centered on lesion load, with noise (to mimic within-subject behavioral variability) added proportional to L × (1 – L), leading to less noise for more extreme lesion loads (near 0 or 1.0). **D.** To simulate behaviors arising from a single Brodmann region, only a single region is selected. **E.** Lesion load is calculated for the single selected region. **F.** Behavior is simulated as in **C**; the only difference between B_1_ and B_2_ is caused by the added noise component.

**Figure S2.**
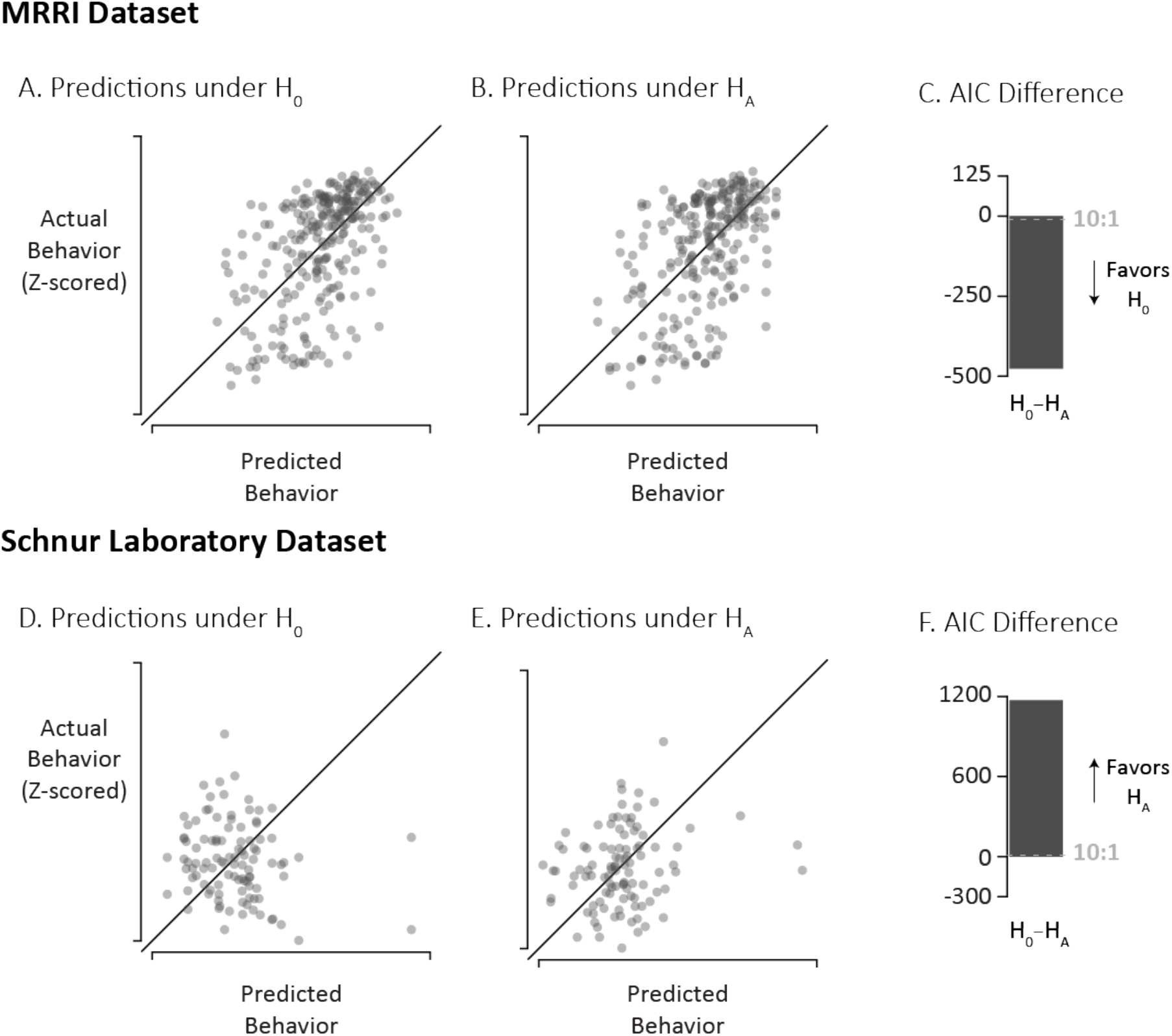
Using PVC with multivariate lesion-behavior maps generated from support vector regression (SVR) provides qualitatively the same results as PVC implemented with SCCAN. Panels A-C. Using the MRRI dataset, the PVC method compares the actual behavior to predictions generated under the null hypothesis (H_0_) and the alternative hypotheses (H_A_). Solid diagonal line indicates perfect prediction. C. The AIC difference was decisive for H_0_ (cutoff at - 10, gray dashed line). Panels D-F are similar to panels A-C except applied to the Schnur Laboratory Dataset, where these data support H_A_: data are predicted by separate LBMs. F. The AIC difference was decisive for H_A_ (cutoff at +10, gray dashed line).

